# Evidence of Incoherent Fluid Flow in the Human Brain from Multidimensional MRI

**DOI:** 10.1101/2025.08.05.668680

**Authors:** Chenyang Li, Yulin Ge, Jiangyang Zhang

## Abstract

**Purpose:** The human brain contains multiple fluids, including blood, cerebrospinal fluid (CSF), and tissue water. While compartmental models have been used to examine microvascular blood perfusion using intravoxel incoherent motion (IVIM), evidence on incoherent flows of CSF is emerging. This study aims to develop in vivo multi-dimensional MRI methods to investigate potential contributions of CSF in the IVIM regime.

**Method:** T_1_-Diffusion (*T_1_-D*) and T_2_-Diffusion (*T_2_-D*) MRI data were acquired from 10 healthy subjects to investigate the relaxivity and diffusion signatures of incoherent fluid flows in the brain. Based on the T_1_-D and T_2_-D results, T_1_/T_2_ selective IVIM protocols were developed to map incoherent CSF flows in the human brains.

**Results:** *T_1_-D* and *T_2_-D* MRI suggested incoherent CSF flow in the human brain subarachnoid space. Results from four different relaxation selective IVIM methods further supported incoherent CSF flows in these regions.

**Conclusion:** We have shown the feasibility of using *T_1_-D* and *T_2_-D* MRI within the low b-value regime to probe the heterogeneity of IVIM components. Designed based on the 2D MRI spectra, relaxation selective 1D IVIM acquisition can be obtained within clinically feasible timeframe.

## 1. Introduction

The human brain contains multiple compartments of various tissues and fluids, ranging from microscopic tissue water and capillaries to large ventricles filled with cerebrospinal fluid (CSF), each characterized by distinct MR parameters (e.g. T_1_/T_2_ relaxivity and diffusivity (*D*))^1^. Since MRI signal is the ensemble average of all tissue and fluid compartments within a voxel, distinguishing signals from individual compartments presents a significant challenge. Successfully separating these compartmental signals offers substantial advantages by enabling the detection of compartment-specific pathology with enhanced sensitivity and specificity.

Several multi-compartment models and related imaging methods have been developed to extract subvoxel distributions of compartments with distinct relaxivity or diffusivity values, as in the case of myelin water imaging^2,3^ and diffusion-based microstructural imaging^4,5^. However, these approaches face inherent limitations when tissue or fluid compartments exhibit similar relaxivities or diffusivities, creating uncertainty in the interpretation of results since such compartments cannot be fully disentangled using single-parameter methods alone. Multidimensional MRI (MD-MRI) offers a powerful framework to address this challenge by jointly encoding relaxivity and diffusivity parameters and uses inverse Laplace transform (ILT) or other methods to reconstruct the joint distribution of these properties among spin populations^6^. This approach enhances the ability to resolve subvoxel compartments that would otherwise remain indistinguishable. For example, recent reports have demonstrated the capability of MD-MRI to map axonal diffuse injury^7^ and astrogliosis in post-mortem human brains^8,9^ associated with traumatic brain injuries and Alzheimer’s disease pathology.

In vivo implementation of MD-MRI, however, remains challenging^10–12^ due to the lengthy acquisition required to cover extensive two-dimensional (2D) spectra via joint relaxation and diffusion encoding. Sparse acquisition schemes, such as compressed sensing^13^, have been proposed to accelerate the 2D acquisition. Recently, Benjamini et al.^14–16^ introduced the marginal distribution constrained optimization (MADCO) technique, which can dramatically accelerate MD-MRI acquisition, potentially for in vivo applications, based on the prior knowledge gained from marginal distribution of relaxivity and diffusivity.

One unique diffusion MRI technique is intravoxel incoherent motion (IVIM) imaging. First introduced by Le Bihan et al.^17–19^, classic IVIM uses a two-compartment model to separate the microvascular compartment, known as pseudo-diffusion, from the tissue water compartment, which has lower diffusivity. Later, an additional compartment with intermediate diffusivities (e.g., the cerebrospinal fluid, or CSF) were added to the IVIM model to account for the free CSF pool^20^. In recent years, several studies have demonstrated the detection of CSF flows using diffusion MRI with low b-values^21–23^, but whether incoherent CSF flows can contribute to the pseudo-diffusion compartment in IVIM imaging has not been fully explored.

In this study, we first implemented in vivo MD-MRI (T_1_-*D*/T_2_-*D*) of the human brain and used the multi-dimensional framework to analyze signals in the IVIM regime. The observed T_1_-*D*/T_2_-*D* spectra indicated potential incoherent flow of the CSF in the subarachnoid space. Furthermore, we have developed relaxation-selective IVIM acquisition methods that can rapidly map this flow in under five minutes by exploiting its unique relaxation and diffusion properties.

## 2. Materials and methods

### 2.1 Human Subjects

The study was approved by the Institutional Review Board at New York University (NYU) Grossman School of Medicine. With HIPAA-compliant and IRB-approved technical development acquisition policy, we recruited ten healthy volunteers (age = 31.1 ± 4.8 years, F/M = 5/5). All participants were given written, informed consent before MRI scans.

### 2.2 T_1_-Diffusion (T_1_-D) and T_2_-Diffusion (T_2_-D) MRI

All MR experiments were performed on a Siemens 3T Prisma system using the standard diffusion-weighted echo planar imaging (EPI) sequence with 64-channel head-coil. Compared to conventional relaxivity and diffusion MRI, MD-MRI jointly encodes both diffusion and relaxation (T_1_ and T_2_). For a typical diffusion MRI experiment, the diffusion signal can be summarized as the weighted summary of diffusion signals of each compartment shown in Eq. [1], where *b* is the diffusion-weighting and *p*(*D*) is the distribution of diffusivity.

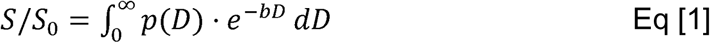

For T_1_-D MRI, an inversion pulse was added before the diffusion encoding (**Fig. 1A**), and the signal (*S*) was modulated by both the inversion time (TI) and *b* as in Eq. [2].

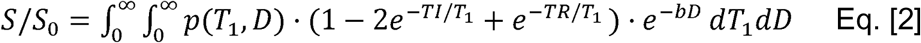

**Figure 1.**
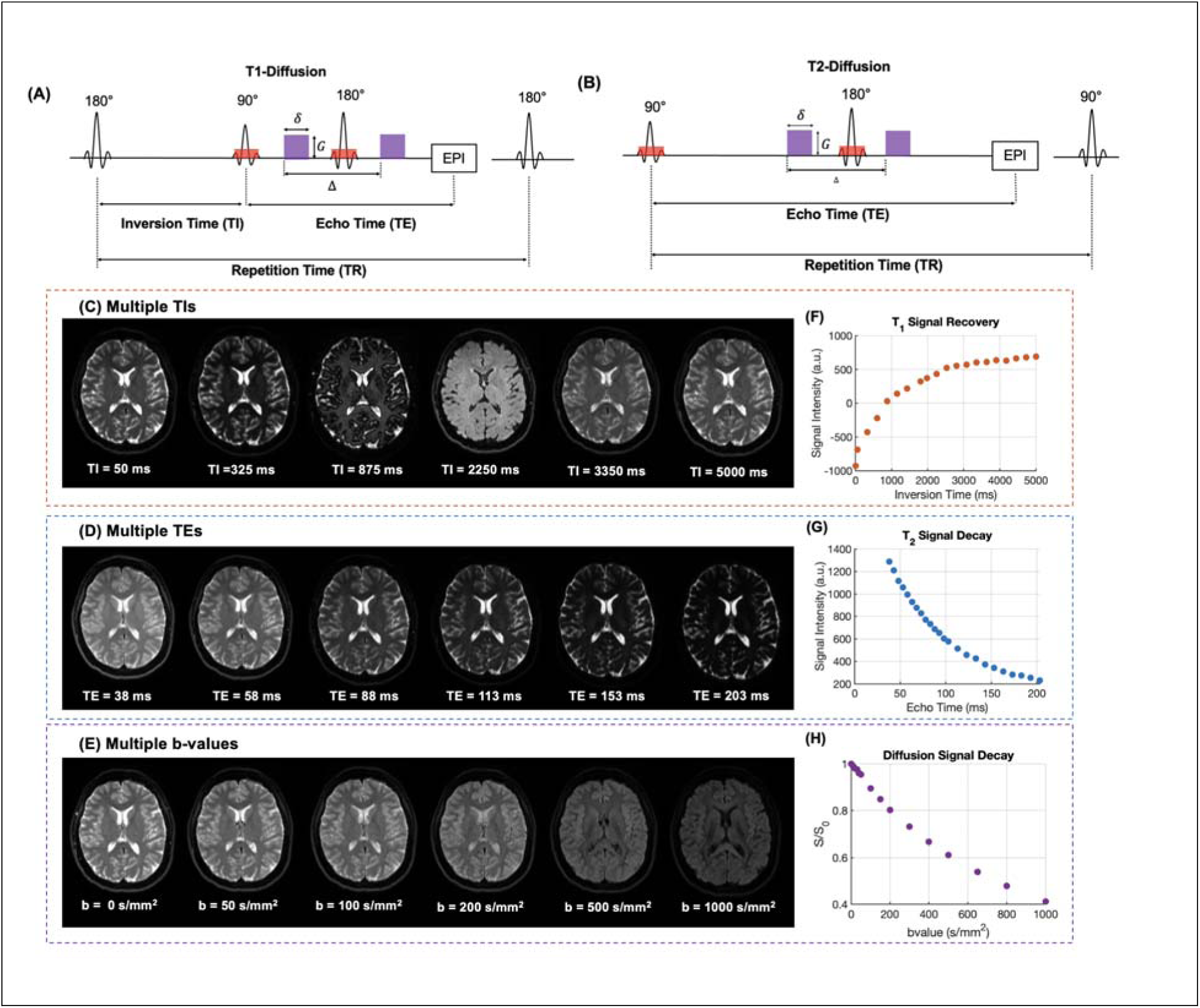
MRI pulse sequences and representative images. (A-B) Pulse sequences for acquiring T_1_-*D* and T_2_-*D* data. (C-E) Representative mTI, mTE, and diffusion-weighted images. (F-H) Representative signals in the cortex showing T_1_ recovery, T_2_ decay, and diffusion decay.

where TE is the echo time and *p(T_1_,D)* is the distributions of protons in the *T_1_-D* domain. For T_2_-D MRI, the signal was modulated by both the TE and diffusion weightings (**Fig. 1B**) as in Eq. [3].

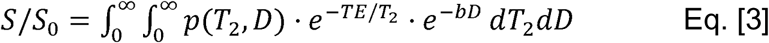

where, and *p(T_2_,D)* is the distribution of protons in the *T_2_-D* domain.

To accelerate the acquisition of T_1_-D and T_2_-D spectra, we used the MADCO acquisition scheme, which needs one-dimensional (1D) marginal distributions *(p(T_1_), p(T_2_),* and *p(D)*) as the prior information. Accordingly, we first acquired non-diffusion-weighting (b_0_) images at 19 TIs (0, 50, 325, 600, 875, 1150, 1430, 1800, 1980, 2250, 2530, 2800, 3080, 3350, 3630, 4180, 4450, 4730, and 5000 ms) with TE/TR = 72/15000ms, and the data were used to calculate the 1D T_1_ spectrum. We then acquired b_0_ images at 24 TEs (38, 43, 48, 53, 58, 63, 68, 73, 78, 83, 88, 93, 98, 103, 113, 123, 133, 143, 153, 163, 173, 183, 193, and 203 ms) with TR = 8000ms, and the data was used to calculate the 1D T_2_ spectrum. We also acquired dMRI data with fifteen b-values (0, 10, 20, 30, 40, 50, 100, 150, 200, 300, 400, 500, 650, 800 and 1000 s/mm^2^) with TE/TR=58/8000ms. These parameters were chosen to capture existing known tissue, blood, and CSF compartments based on their values in the literature. We acquired 55 axial slices with a field of view of 288 mm x 288 mm, a slice thickness of 3 mm, a matrix size of 192 x 192, and the native resolution of 1.5 x 1.5 x 3 mm^3^. The acquisition time of the T_1_, T_2_, and diffusion scans were 14 and 15, and 3 minutes, respectively. Representative images and signal attenuation curves are shown in **Fig. 1C-H**.

Following the acquisition of the 1D spectra, sparsely sampled IVIM data at five TEs (38, 58, 78, 153, and 183 ms), seven TIs (0, 50, 325, 600, 875, 1800, and 3900 ms), and fifteen b-values (0, 10, 20, 30, 40, 50, 100, 150, 200, 300, 400, 500, 650, 800, and 1000 s/mm^2^) were acquired. All 1D and 2D T_1_-D and T_2_-D MRI data were co-registered. The acquisition time of the 5 x 7 x 15 2D T_1_-T_2_-D data was approximately 30 minutes. Together with the 1D acquisitions, the total acquisition time of T_1_-D and T_2_-D data was around 1 hour and 5 minutes.

### 2.3 Image processing and MADCO reconstruction

The MADCO data was preprocessed using the DESIGNER pipeline^24^, which includes distortion correction, denoise and bias field correction. The voxel-wise marginal distribution of T_1_, T_2_ and diffusivity were then calculated using 1D inverse Laplace transform (ILT) based on L_2_ regularization and used as prior knowledge for the MADCO reconstruction^25^. Similar to the T_1_-D spectral analysis by Avaram et al.^10^, the polarity of T_1_-D data was adjusted for phase correction. The diffusion signals with different TE were used to solve the joint distribution of relaxation-diffusion spectrum using the MADCO framework and reconstructed using the method as described by Benjamini et al^14,15^ and Zong et al^11^.

### 2.4 T_1_-D and T_2_-D MRI informed FLAIR-IVIM, LongTE-IVIM and VASO-LongTE-IVIM

In this study, we performed three types of T_1_- and T_2_-selective IVIM acquisitions and compared the results with conventional IVIM. All 1D IVIM data were acquired using a diffusion-weighted EPI sequence with fifteen b–values (0, 10, 20, 30, 40, 50, 100, 150, 200, 300, 400, 500, 650, 800 and 1000 s/mm^2^), the single diffusion direction along the slice direction, and the same field-of-view and matrix size as the T_1_-D/T_2_-D experiments. We previously used these relaxation selective IVIM acquisitions to study fluid compartments in the choroid plexus^26^.

1. Conventional IVIM (**Fig. 2B**) scans were performed with TE/TI/TR of 73/0/8000ms. Here, zero TI indicates no inversion pulse before diffusion encoding.
2. FLAIR-IVIM (**Fig. 2C**) scans were performed with TE/TI/TR=73ms/1800ms/8000ms to effectively suppress CSF signals.
3. LongTE-IVIM (**Fig. 2D**) scans were performed with TE/TI/TR=183ms/0ms/8000ms to suppress the tissue signal while preserving blood and CSF signals due to their relatively longer T_2_ relaxation time.
4. Vascular space occupancy (VASO)-LongTE-IVIM (**Fig. 2E**) scans were performed with TE/TI/TR=183ms/1150ms/8000ms. The TI was chosen to suppress blood signals based on previously reported blood T_1_ values at 3T^27^ (T_1,blood_ = 1624ms), and the long TE was used to suppress tissue signals. Theoretically, under this IVIM acquisition, only the CSF signal remained, which facilitated the identification of incoherent CSF IVIM components.

**Figure 2.**
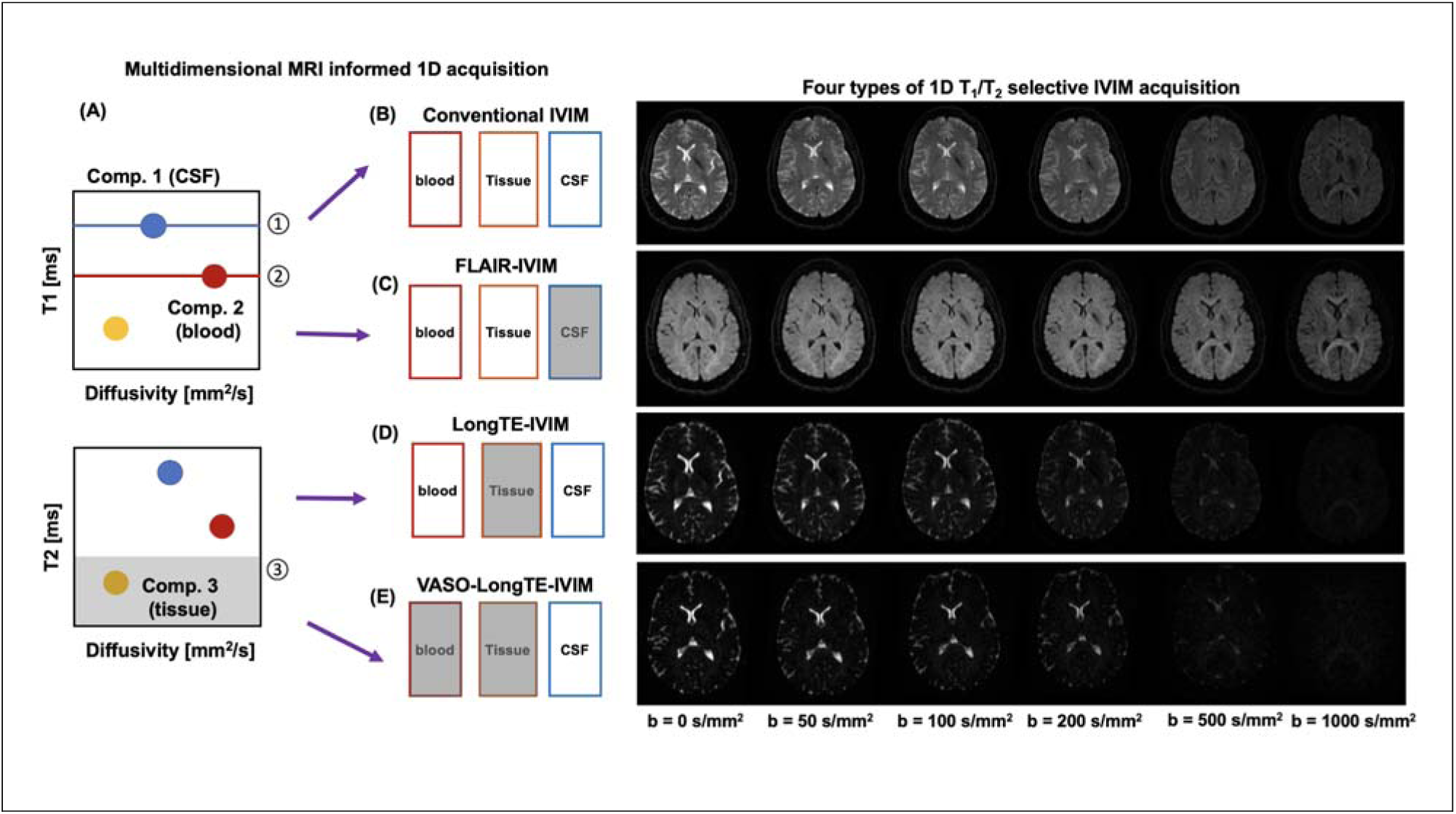
Relaxation selective IVIM methods. (A) A schematic representation of the four IVIM acquisition methods designed to suppress fluid compartments based on T_1_-*D* and T_2_-*D* 2D spectra. (B) In conventional IVIM, signals contain contributions from blood, tissue, and CSF. (C) FLAIR-IVIM suppresses CSF signal through inversion recovery preparation; (D) LongTE-IVIM attenuates tissue signals with relatively short T_2_ values; (E) VASO-LongTE-IVIM combines blood nulling and long echo time to suppress both blood and tissue signals.

Each sequence took approximately 2 minutes and 50 seconds. To account for T_1_ and T_2_ effects induced by T1 preparation and different TEs, the diffusion data were normalized with their individual S_0_ values. To analyze the diffusion spectrum, 1D ILT was applied on the IVIM data to derive the distributions of diffusivity values. Following Wong et al.^28^, a peak window with 10^−4^ to 1.5×10^−3^ mm^2^/s was used for the parenchymal tissue, and the area under the curve within this window divided by the total area was the volume fraction of tissue water diffusion (f_tissue_). Peak window from 1.5×10^−3^ to 4×10^−3^ mm^2^/s and from 4×10^−3^ to 1 mm^2^/s were used for free diffusion (mainly CSF) and pseudo-diffusion, respectively, and volume fractions f_free_ and f_ivim_ were calculated in the same way as f_tissue_.

## 3. Results

### 3.1 The 1D marginal distributions of T1, T2, and diffusivity in the human brain

As expected, the 1D marginal distributions were regional-dependent. In the ventricle, the 1D distribution showed a single peak, indicating a spin population, with T_1_ and T_2_ of CSF and diffusivity of free water (**Fig. 3**, left column). In the subarachnoid space, the 1D distribution of T_1_, T_2_, and diffusivity distributions showed additional peaks with lower T_1_, T_2_ and higher diffusivity values than CSF, indicating additional fluid compartments (**Fig. 3**, middle column). In the cerebral cortex, the distributions also indicated several fluid compartments (**Fig. 3** right column). Interestingly, the 1D distribution of diffusivity in the subarachnoid space also showed a peak with diffusivity in the IVIM regime (diffusivity higher than free diffusion), similar to the IVIM peak observed in the cortex. The relationships among the peaks observed in the three 1D distributions remained unclear. For example, whether the IVIM peak in the subarachnoid space was from single or multiple fluids was difficult to be determined based solely on 1D data.

**Figure 3.**
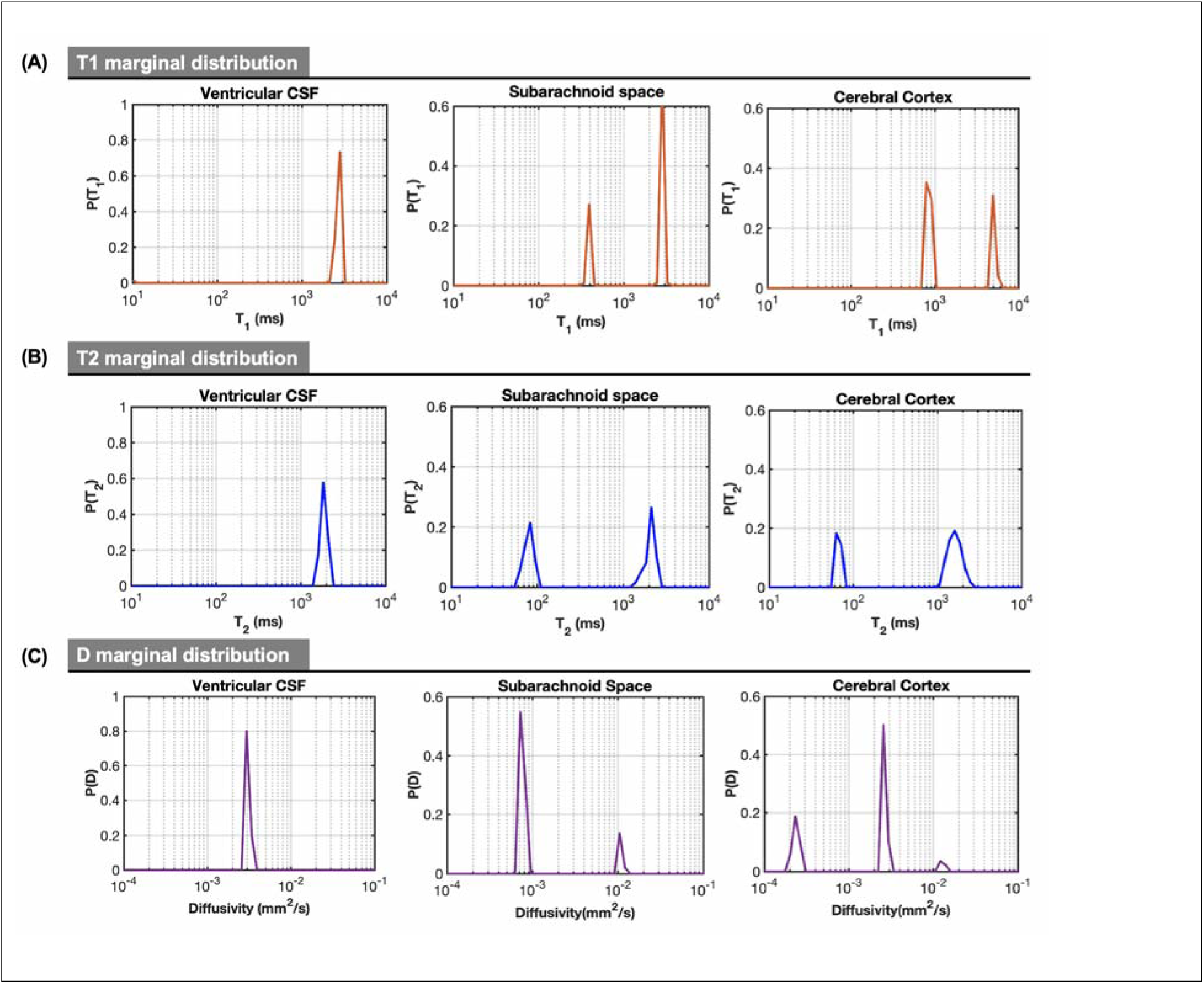
Representative T_1_, T_2_, and diffusivity distributions in the ventricle, subaranchoid space, and cortex. The measurements from the ventricles were used as references for CSF in other regions.

### 3.2 In vivo T_1_-D and T_2_-D spectra in the IVIM regime

T_1_-D and T_2_-D data provided additional insights into the fluid composition of the IVIM peak in the SAS space. In the T_2_-D data (**Fig. 4**), two peaks in the IVIM region were observed with T_2_ values close to blood and CSF, potentially suggesting incoherent CSF flow that resides in the IVIM range. The T_1_-D data, however, showed a strong IVIM peak with T_1_ values close to CSF. These results suggested the existence of incoherent CSF flows that contributed to the IVIM signals.

**Figure 4.**
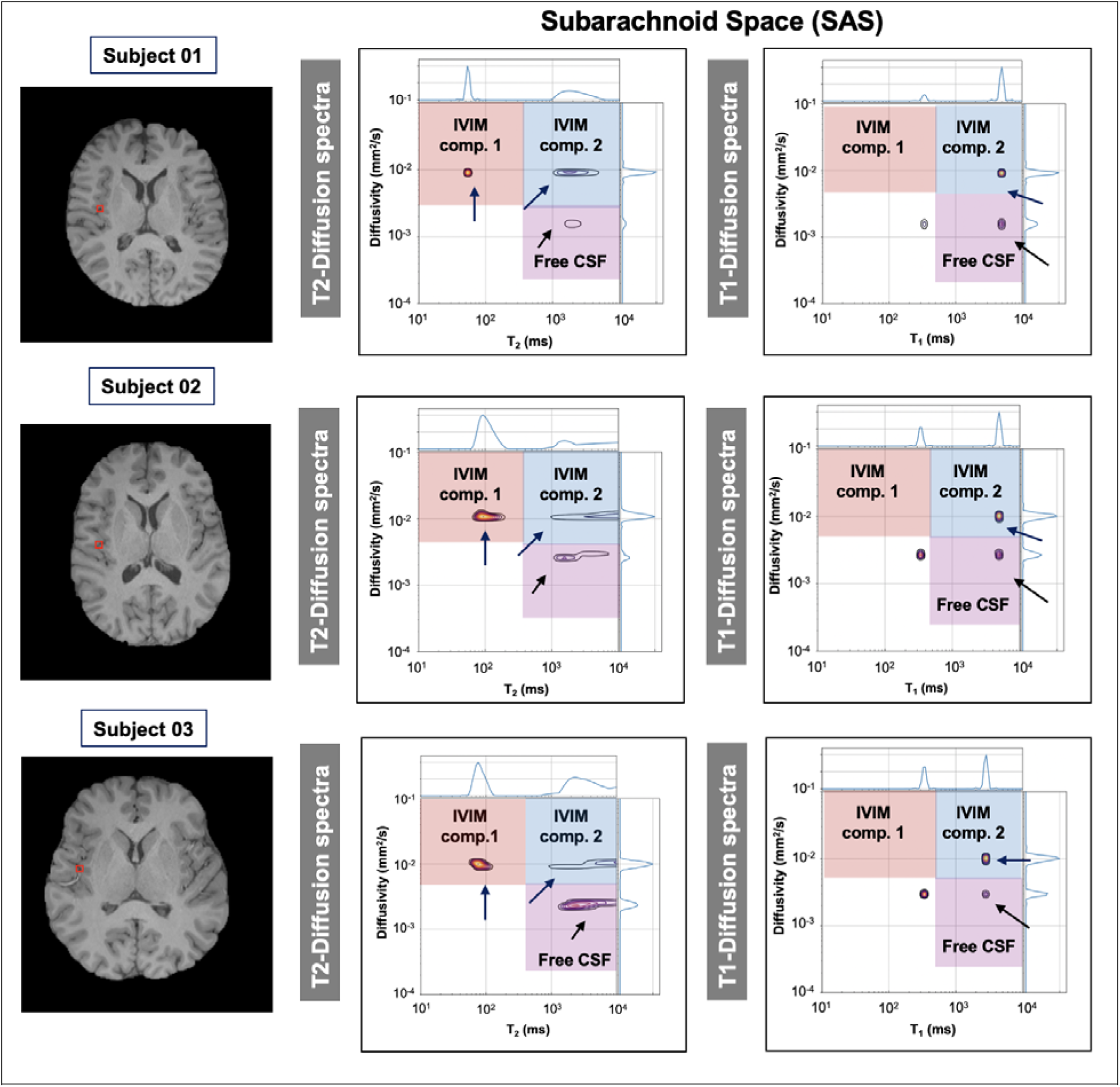
Representative SAS T_1_-*D* (A) and T_2_-D (B) correlation spectra from three subjects, demonstrating two IVIM components (within the red and blue rectangles, respectively), and one CSF component (within the purple rectangle).

### 3.3 Further investigation of the potential incoherent CSF flow using relaxation selective IVIM

Although the 2D T_1_-D and T_2_-D acquisitions provided key information, the time-consuming acquisition limits their clinical applications. The 2D findings guided the design of more targeted 1D experiments with shorter acquisition time to examine potential incoherent CSF flow. Conventional IVIM signals from the ventricles and cortex showed single and three peaks, whereas the signals from the SAS showed an IVIM peak and a free-diffusion peak (**Fig. 5A**). FLAIR preparation effectively suppressed CSF signals, resulting in no visible signals in the ventricles and SAS while preserving three peaks in the cortex (IVIM, tissue, and free diffusion, likely from CSF) (**Fig. 5B**). Interestingly, the presence of the free diffusion peak after FLAIR preparation suggests free water compartments with T_1_ values distinct from the CSF. Long TE sequences designed to suppress tissue signals substantially attenuated cortical signals yet maintained strong IVIM and free diffusion peaks in the SAS (**Fig. 5C**). Adding VASO preparation, which suppresses blood signals, did not completely remove the IVIM peak (**Fig. 5D**). Voxel-wise analysis confirmed that LongTE-IVIM and VASO-LongTE-IVIM components were predominantly localized to the SAS (**Fig. 6**), with corresponding volume fractions and diffusivities of the free and IVIM components detailed in **Table 1**.

**Figure 5.**
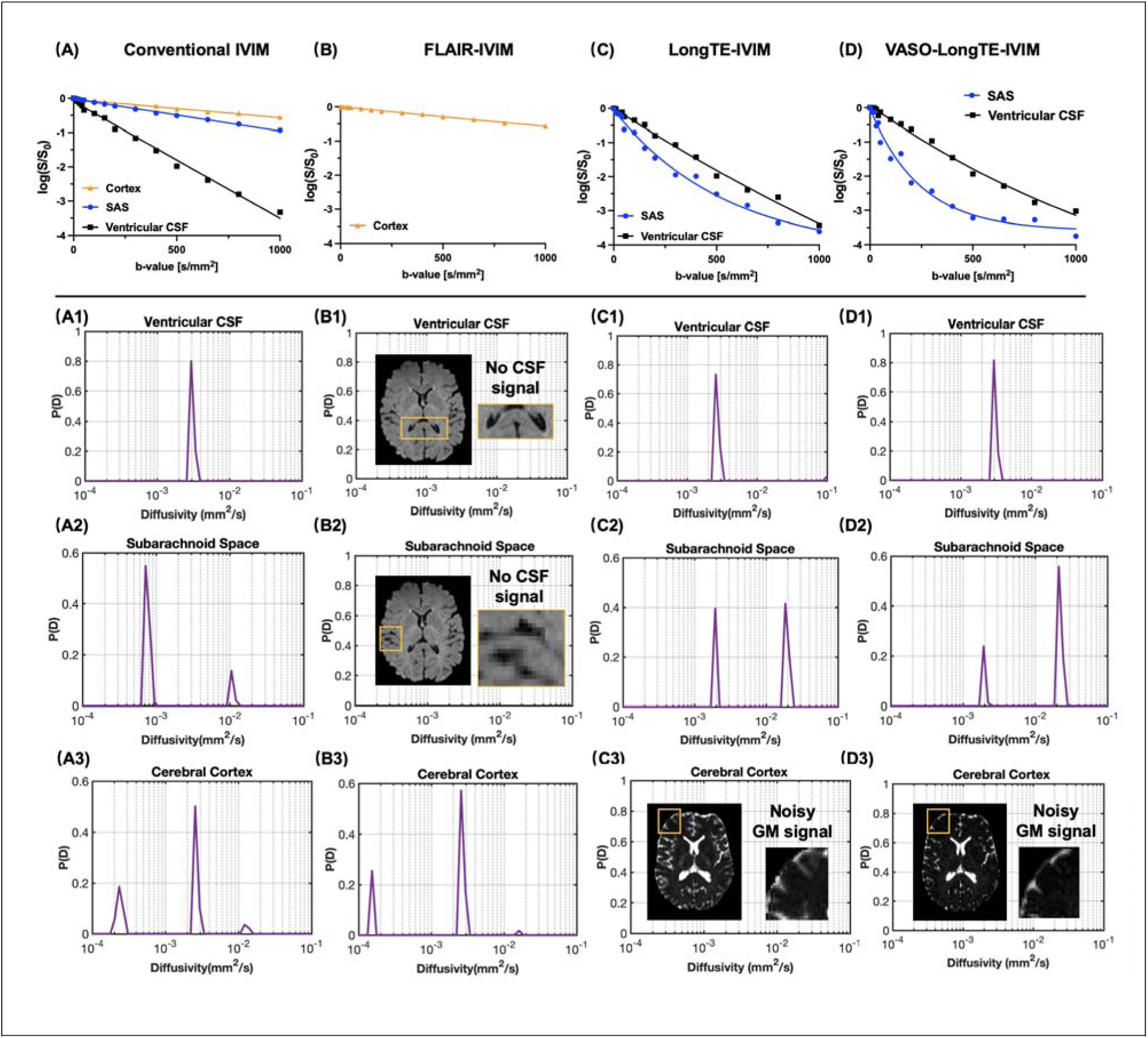
Representative log-scale signal decay from four IVIM acquisition strategies:(A) Conventional IVIM, (B) FLAIR-IVIM, (C) LongTE-IVIM, and (D) VASO-LongTE-IVIM in Cortex, SAS and ventricular CSF. Corresponding 1D inverse Laplace transform (ILT) results showing the distribution of diffusivities (P(D)) in ventricular CSF (A1-D1), SAS (A2-D2) and Cortex (A3-D3). Ventricular CSF shows mono-exponential decay with a single diffusion peak; SAS and cortex exhibit multi-component diffusion profiles. Note that FLAIR-IVIM suppresses ventricular CSF signal, and no physically meaningful diffusion profile is obtained for ventricular CSF in FLAIR-IVIM acquisition.

**Figure 6.**
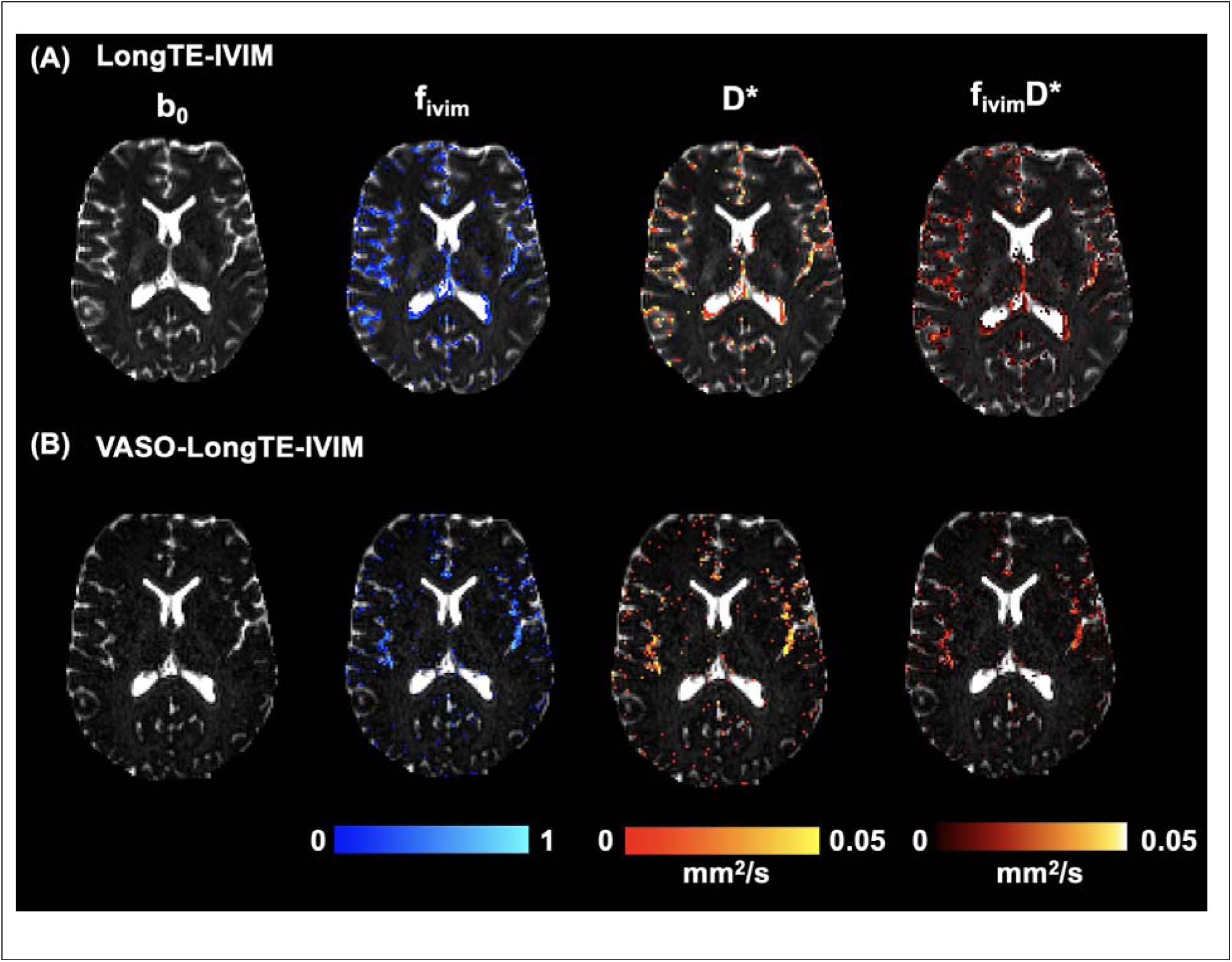
Voxel-wise ILT mapping of (A) LongTE-IVIM and (B) VASO-LongTE-IVIM highlights that the f_IVIM_ components in the brain were mostly located near cerebral cortex and subarachnoid space.

**Table 1.** Reported volume fraction and diffusivity of two IVIM components in the SAS space characterized by T2-Diffusion MRI spectra. Mean values and standard deviations from 10 healthy subjects are shown here.

| Subarachnoid space (SAS) | $f_{\text{free}}(D)$ | $D_{\text{free}}$<br>($\times 10^{-3} \text{ mm}^2/\text{s}$) | $f_{\text{IVIM}}(D)$ | $D_{\text{IVIM}}$<br>($\times 10^{-3} \text{ mm}^2/\text{s}$) |
| --- | --- | --- | --- | --- |
| LongTE-IVIM | $0.43 \pm 0.13$ | $3.1 \pm 0.9$ | $0.55 \pm 0.17$ | $39.13 \pm 14.5$ |
| VASO-LongTE-IVIM | $0.35 \pm 0.04$ | $2.9 \pm 0.6$ | $0.61 \pm 0.05$ | $30.8 \pm 16.4$ |

## 4. Discussion

In this study, we performed in vivo MD-MRI and used the MD-MRI framework to explore the heterogeneity of IVIM components in the human brain and developed 1D relaxation selective IVIM techniques based on the 2D *T_1_-D* and *T_2_-D* spectra.

### 4.1 Diffusion MRI of CSF flows

CSF flow is gaining attention as the primary driving force for waste clearance in the brains, making techniques that can measure CSF flow increasingly important. As described by Wright et al.^29^, CSF movement in the human brain comprises both coherent and incoherent flow components, reflecting the complex dynamics of CSF circulation. Coherent flow typically arises from bulk CSF movement driven by arterial pulsation or respiration and has been well characterized in previous studies using velocity-encoded MRI, which demonstrated relatively fast, directional CSF flow^30–32^. In contrast, incoherent flow refers to more randomized and multidirectional, potentially resulting from diffusion-like displacement. The subarachnoid space (SAS) contains a complex network of trabeculae that creates a spider-web-like microenvironment, potentially leading to incoherent fluid flow patterns^33,34^.

Emerging evidence has suggested the feasibility of using diffusion MRI to detect CSF flow. Becker et al.^35^ reported that a two exponential model can capture CSF flows in the ventricles. Later, Wen et al.^36^ and Han et al.^37^ have used a dynamic diffusion-weighted imaging technique to suppress the fast blood flow and observed the dynamic ADC measures to characterize CSF flow with cardiac cycle, which is deemed driven primarily by arterial pulsation or respiratory. More recently, Chen et al.^38^ assessed the contribution of CSF flows to D*, including in the SAS, indicating the potential of IVIM to detect incoherent CSF flow.

However, conventional IVIM faces several key limitations in mapping CSF flow: (1) IVIM is not sensitive to fast, coherent CSF movements in the ventricles; (2) difficulty separating incoherent CSF flow from blood flow, as both exhibit diffusivities higher than free diffusion. The MD-MRI framework can potentially address the second limitation by separating blood and CSF flows based on their distinct relaxation properties ^39^.

### 4.2 In vivo MD-MRI of the human brain

While MD-MRI provides rich information, it requires lengthy acquisition times to collect data with multiple diffusion and relaxation weightings. This constraint has led many researchers to conduct MD-MRI studies using in vitro or ex vivo specimens, although in vivo applications are emerging with sparse sampling methods such as compressed sensing^13^ and MADCO^14,15^. Recent advances include Avaram et al.^10^ reporting the first T_1_-D correlation of the in vivo human brain, Slator et al.^40^ combing T * relaxation and IVIM to study the placenta oxygen and perfusion measures, and Martin et al.^41^ and Johnson et al.^12^ proposing higher dimensional frameworks to study the correlation of diffusion frequency, tensor shape with relaxation properties.

The above-mentioned studies primarily acquired data with medium-to-high b-values to disentangle sub-voxel tissue compartments, while MD-MRI with low b-values or IVIM scheme remains unexplored, because previous MD-MRI studies focused on tissue microstructure rather than flow dynamics. Implementing in vivo MD-MRI within the IVIM regime provides a new approach to disentangle contributions from different fluid components to IVIM signals. The T_1_-D and T_2_-D spectra in the IVIM regime provide a comprehensive view of potential sub-voxel IVIM-type flow distributions, enabling the development of faster, targeted acquisitions to validate MD-MRI results by using spectral signatures to isolate specific IVIM signals through relaxation selection in 1D IVIM acquisitions.

Despite using MADCO to accelerate image acquisition, the total imaging time remained substantial in this study. Consequently, only one diffusion encoding direction was used in this study, limiting our ability to capture potential anisotropic CSF flows in the white matter. Additionally, the selection of inversion time in the T_1_-D experiments may affect the resulting spectrum. For voxels that contain multiple fluid compartments, it is difficult to determine the zero-crossing points, resulting in the so-called ‘bounce point artifact’. This may explain that only CSF flow peak was observed in the T_1_-D experiments while both CSF and blood flow peaks were found in the T_2_-D experiments. The T_1_-D spectrum could be improved by avoiding problematic TIs during MADCO reconstruction, with simulations helping to confirm optimal TIs within this range. Furthermore, while ILT offers advantages—requiring minimal assumptions about spin population numbers and MR properties—it also presents disadvantages. ILT is ill-posed and noise-sensitive, with existing applications relying on fine-tuned regularization factors to suppress spurious results. Therefore, the T_1_-D and T_2_-D spectra observed here require further validation.

### 4.3 T1-D and T2-D MRI informed faster 1D relaxation selective acquisition

Although several studies have used relaxation properties to modulate the sensitivity and specificity of diffusion MRI signal, for example, De Santis et al.^42^ have utilized the difference of T_1_ relaxation time to resolve the crossing fibers for relaxivity estimation. Wong et al.^28^ previously employed inversion recovery preparation in IVIM signal to suppress CSF signals, and subsequent ILT analysis on this IR-IVIM signal revealed residual unsuppressed free-diffusion spins in the white matter hyperintensities in small vessel disease patients. Wong et al.^28^ hypothesized this spin population to be interstitial fluid (ISF), where the different fluid components may alter the T_1_ relaxation time that is different from free CSF. Using the T_1_-D or T_2_-D spectra, a more specific acquisition design is possible within clinically feasible timeframe, and a generalized framework extending from T_1_-D and T_2_-D to 1D relaxation-selective diffusion imaging is promising. A previous study by Li et al.^26^ implemented FLAIR-IVIM and VASO-LongTE-IVIM to investigate the fluid compartments within the human choroid plexus. By suppressing surrounding CSF using FLAIR inversion and blood signals using VASO, the complexity of the IVIM signal may be reduced through selective nulling of diffusion components based on the distinct relaxation times of blood and CSF.

Using SAS near the cortex as a representative example, we have found the presence of remaining IVIM components in both LongTE-IVIM and VASO-LongTE-IVIM. Although it is difficult to separate the blood and CSF from the IVIM peaks in LongTE-IVIM due to the relatively longer T_2_ of both blood and CSF. VASO-LongTE-IVIM provides evidence of the existence of CSF flow when blood signal is nulled. The voxel-wise mapping of f_IVIM_ of LongTE-IVIM and VASO-LongTE-IVIM revealed the spatial distribution of IVIM components, which are located mainly in the SAS space. This combination serves as a new imaging approach for detecting incoherent CSF flow in the SAS, showing faster acquisition times and greater potential for clinical translation and application. Similarly, we also expected incoherent CSF flow in the cortex, but capturing it remains challenging due to small volume fractions (**Fig. 5**). According to Saib et al.^43^, using a heavily T_2_-weighted MRI, it showed that the perivascular space in the cortex is within the range of nanoliter scale. Another caveat is that the suppression of blood signals using VASO preparation depends on hematocrit and oxygenation levels^44^, and partial suppression of blood signals will result in residual blood IVIM signals, leading to over-estimation of incoherent CSF flow.

### 4.4 Other technical considerations of in vivo MD-MRI

The parameters used in this study, such as TE and b-values, were designed only for detecting selected groups of spin populations, not all possible spin populations in the brain. For instance, the myelin water in white matter, which has a much shorter T_2_ than tissue, is not detectable in our MD-MRI experiments due to the relatively long TEs (shortest TE of 58ms). The intra-axonal compartment is another tissue compartment that is not detected in this study due to its low diffusivities, which will require stronger diffusion weightings (e.g., b values > 2,000 s/mm^2^). These compromises were made to acquire the 2D spectra within a reasonable time and due to our focus on the flow. Lastly, the diffusion time often changes with varying TEs with clinical diffusion MRI sequence. Previous evidences suggest that IVIM signals may change with diffusion time in the mouse brain and prostate^45^, the time dependent IVIM relies on the architecture of the vessel network, which complicates the interpretation of *T_2_-D* results by including both microstructure, flow and exchange effects. Future works by including diffusion time as another dimension is needed to better understand the effects of microvascular networks and their corresponding spin migration.

## 5. Conclusion

In this study, we demonstrated a proof-of-concept of in vivo MD-IVIM imaging of the human brain, which revealed potentially more than one IVIM component with distinct T_1_-D and T_2_-D signatures. The MD-IVIM results were supported by T_1_ and T_2_ selective relaxation IVIM, suggesting the existence of incoherent CSF flow in the subarachnoid space.

